# Highly variable COI haplotype diversity between three species of invasive pest fruit fly reflects remarkably incongruent demographic histories

**DOI:** 10.1101/742007

**Authors:** Camiel Doorenweerd, Michael San Jose, Norman Barr, Luc Leblanc, Daniel Rubinoff

**Affiliations:** University of Hawaii, Department of Plant and Environmental Protection Services, 3050 Maile Way, Honolulu, Hawaii, 96822-2231, United States; Center for Plant Health Science and Technology, Mission Laboratory, USDA-APHIS, Moore Air Base, 22675 North Moorefield Rd., Ediburg, TX 78541, United States; University of Idaho, Department of Entomology, Plant Pathology and Nematology, 875 Perimeter Drive, MS2329, Moscow, Idaho, 83844-2329, United States

**Keywords:** peach fruit fly, guava fruit fly, melon fly, biosecurity, cox1, dna barcoding, Tephritidae, Dacini, *Bactrocera*, *Zeugodacus*

## Abstract

Distance decay principles predict that species with larger geographic ranges would have greater intraspecific genetic diversity than more restricted species. However, invasive pest species may not follow this prediction, with confounding implications for tracking phenomena including original ranges, invasion pathways and source populations. We sequenced an 815 base-pair section of the COI gene for 441 specimens of *Bactrocera correcta*, 214 *B. zonata* and 372 *Zeugodacus cucurbitae*; three invasive pest fruit fly species with overlapping hostplants. For each species, we explored how many individuals would need to be included in a study to sample the majority of their haplotype diversity. We also tested for phylogeographic signal and used demographic estimators as a proxy for invasion potency. We find contrasting patterns of haplotype diversity amongst the species, where *B. zonata* has the highest diversity but most haplotypes were represented by singletons; *B. correcta* has ∼7 dominant haplotypes more evenly distributed; *Z. cucurbitae* has a single dominant haplotype with closely related singletons in a ‘star-shape’ surrounding it. We discuss how these differing patterns relate to their invasion histories. None of the species showed meaningful phylogeographic patterns, possibly due to gene-flow between areas across their distributions, obscuring or eliminating substructuring.

## Introduction

A fundamental assumption of species-level research is that sampling specimens from across their range accurately represents intraspecific genetic diversity (Steinke et al. 2009; Bergsten et al. 2012; Fernandez-Triana et al. 2014; Huemer et al. 2018). This tenet is based on population-level processes such as genetic drift and selection, which follow the distance decay principle (Nekola and White 1999) resulting in regional differences that increase with geographic distance as the probability of gene-flow between more distant populations decreases. These divergences may be augmented in heterogenous landscapes or under strong environmental gradients (Alonso-Blanco et al. 2016). Indeed, studies using various, predominantly native, taxa over large geographic ranges typically find that species with broader distributions, on average, have more intraspecific diversity than species with more restricted distributions (e.g. Lukhtanov et al. 2009; Bergsten et al. 2012; Huemer et al. 2018). However, depending on the life-history characters and evolutionary history, intraspecific genetic diversity can be regionally heterogeneous and may vary widely between taxa (e.g. Alonso-Blanco et al. 2016; Domyan and Shapiro 2017).

We tested distance decay assumptions with three of the worst agricultural pests in Southeast Asia, since this has implications for their control, quarantine and the reconstruction of invasion patterns based on demographic history. When intraspecific variation is correlated with geography it becomes possible to trace the geographic origins of invasive species and provides a framework to prevent future incursions and better manage invasive populations (Armstrong and Ball 2005; Zhang et al. 2013; Dupuis et al. 2018b). Moreover, populations can be adapted to their local environments, which influences the chance of a successful future invasion (Violle et al. 2014) or can even reinvigorate invasions by enabling access to new hosts or through biocontrol resistance (Mimura et al. 2017; Reil et al. 2018). If, on the other hand, no intraspecific variation is detected in the invasive range, it suggests a single small initial invasive population or multiple invasions from the same source. In this situation, finding the source of the invasion can be important in predicting infestation potential and to locate natural enemies for control. This kind of information would be important for understanding the invasion threat posed by other serious pests, an increasingly common problem as globalization intensifies.

When using molecular methods for species identification, commonly using the Cytochrome C Oxidase I (COI) gene, most automated species delimitation algorithms, such as BIN, CROP, jMOTU, GMYC and ABGD (for comparisons see (Ratnasingham and Hebert 2013; Pentinsaari et al. 2014; Kekkonen et al. 2015), rely, at least in part, on there being a diagnosable difference between intra- and interspecific variation and then averaging these values within phylogenetic groupings, in order to ultimately detect differences between species. Commonly, large DNA barcoding initiatives have focused on sequencing as many species as possible, with the implicit assumption that genetic diversity has been adequately sampled to reflect species identity (Funk and Omland 2003; Ward et al. 2005; Kerr et al. 2007; Steinke et al. 2009; Mutanen et al. 2016; Gibbs 2017). Few studies have focused on haplotype accumulation, and those that have studied it found that even with sample sizes >50, generally only a small portion of the actual diversity was sampled (e.g. Bergsten et al. 2012; Phillips et al. 2015). Little attention has been paid to what qualifies as sufficient sampling for reliable delimitation, identification or population assignment, with potentially significant ramifications for monitoring pest species in the context of control and quarantine.

Larvae of the globally distributed fruit fly family Tephritidae feed on fruit or flowers and roughly 250 of the ca. 5,000 species have become agricultural pests, some of which have greatly expanded their distributions due to human activity (White and Elson-Harris 1992; Norrbom et al. 1998; Meyer et al. 2007; Vargas et al. 2015). The recent demographic and geographic changes of pests in this group make them ideal models for testing distance decay predictions, the results of which would be directly relevant to other invasive pest systems. The billions in USD of economic damage that tephritid fruit flies inflict across the planet (Siebert and Cooper 1995; Badii et al. 2015; CNAS 2015) has sparked multiple initiatives to build COI DNA barcode reference libraries in order to detect and identify problematic species (Armstrong and Ball 2005; Virgilio et al. 2012; Smit et al. 2013; Jiang et al. 2014; Aketarawong et al. 2015; Barr et al. 2017). Three invasive polyphagous pests; peach fruit fly - *Bactrocera zonata* Saunders, guava fruit fly - *Bactrocera correcta* Bezzi and melon fly - *Zeugodacus cucurbitae* Coquillett have overlapping host ranges that include over fifty species of commercially grown fruit, as well as partly overlapping distribution ranges (Allwood et al. 1999; Vargas et al. 2015; Fig. 1). *Zeugodacus cucurbitae* is the most invasive of the three, and has spread out of South-East Asia across Africa and much of the Pacific (De Meyer et al. 2015). *Bactrocera zonata* is likely native to the Indian subcontinent and is spreading westward into northern Africa. The extent of the invasive versus the native ranges of *B. correcta* have not been studied in much detail, but likely include parts of South-East Asia. Plant protection agencies, including the United States Department of Agriculture and the European Plant Protection Agency have dedicated programs to exclude the introduction and establishment of these insects (CABI 2018; EPPO 2018). However, none of these species have hitherto been studied molecularly throughout their range.

**Figure 1.**
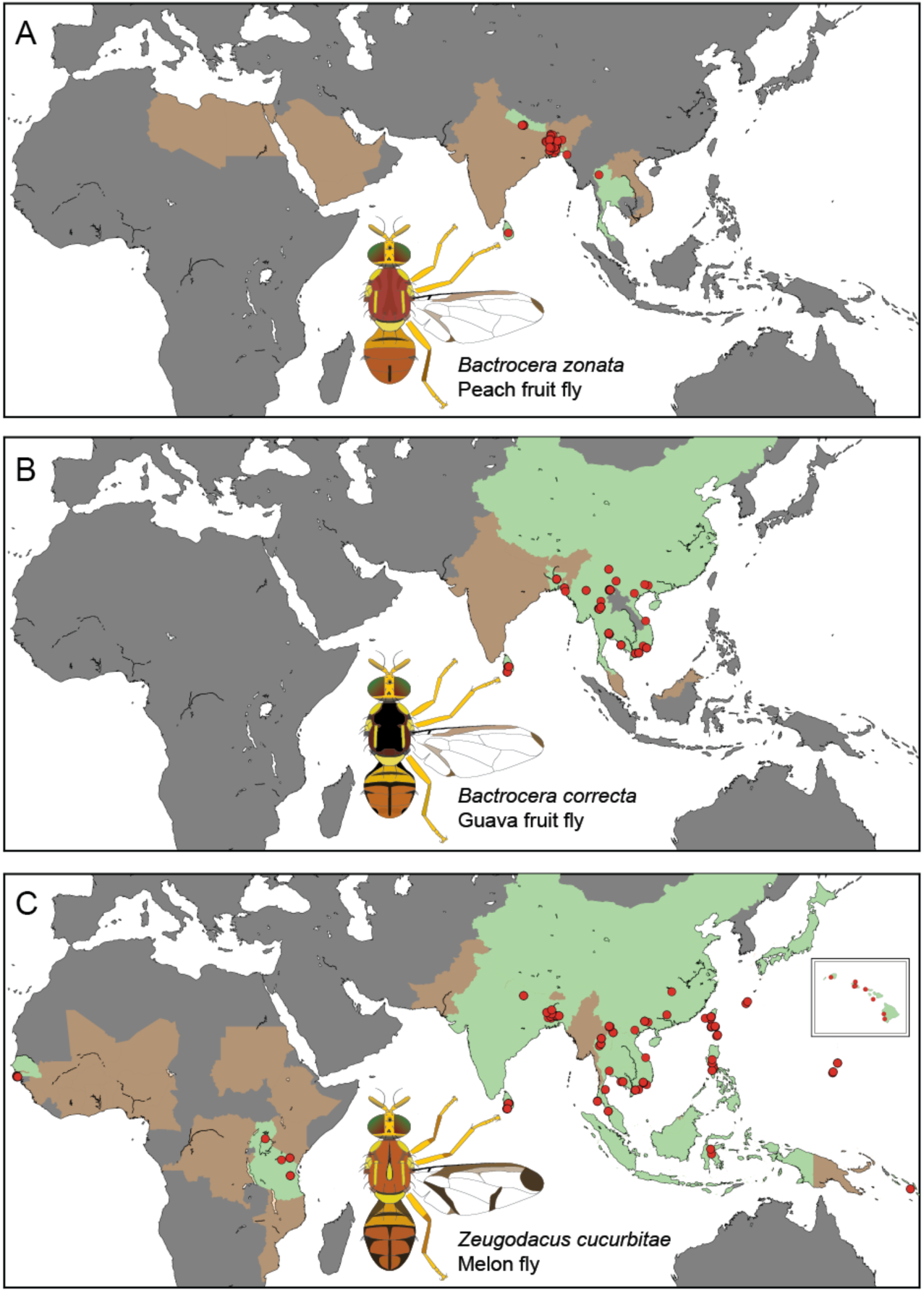
Distribution and sampling points of the three species; A. *Bactrocera zonata*, B. *Bactrocera correcta* and C. *Zeugodacus cucurbitae*. Green areas indicate countries for which the species has been reported and we have sampled, brown areas indicate countries for which the species has been reported but we were unable to sample. Red circles indicate sampling localities. The map insert in C shows the Hawaiian Islands. Note that the actual distributions of the species do not always follow political boundaries, the flies do not occur in northern China, or the northern Islands of Japan.

We performed high-density sampling of *Zeugodacus cucurbitae* and *Bactrocera correcta* and *B. zonata*, whereby we attempted to include samples from across their ranges. Using an 815 base-pair section of the COI gene (3’-P region) we explored with rarefaction curves how many specimens hypothetically needed to be sampled to obtain all the haplotypes. We additionally tested whether the species could be identified reliably using COI and looked for a phylogeographic signal that may be useful in quarantine efforts and for reconstructing invasion pathways. Finally, we evaluated the efficacy of demographic estimators to infer historic and current population expansion as a proxy for pest severity and future invasion threat.

## Material & Methods

### Taxon sampling

Based on morphology and a seven-gene molecular phylogeny with 178 Dacini species (San Jose et al. 2018a), *B. correcta* and *B. zonata* are sister species, and *Z. cucurbitae* is closely related to *Z. tau* Walker, *Z. synnephes* Hendel and *Z. choristus* May. *Zeugodacus cucurbitae* is widespread and is documented to be expanding its distribution, which currently includes much of subtropical Africa, South-East Asia and Pacific Islands, including Guam and Hawaii (De Meyer et al. 2015; Vargas et al. 2015), Fig. 1C). *Bactrocera zonata* is present in northern Africa and Saudi Arabia, the Indian subcontinent and has been reported from Thailand and Vietnam (Fig. 1A). Partly overlapping with this distribution, *B. correcta* can be found throughout most of mainland South-East Asia (Fig. 1B) (Drew and Romig 2013). Samples were collected between 2005 and 2017 (Fig. 1, BOLD doi: xxx-xxx-xxx). *Bactrocera correcta* and *B. zonata* were collected with the male attractant methyl eugenol, and *Z. cucurbitae* with cue lure (Metcalf et al. 1983). Bucket traps with one of these lures and a 1×1cm dichlorvos strip as a killing agent were maintained for three to five days along trails in natural areas or near agricultural land. Specimens were preserved in 95% ethanol and stored at −20°C. We selected 441 *B. correcta*, 214 *B. zonata* and 372 specimens of *Z. cucurbitae* for DNA extraction. All identifications were done based on morphology.

### DNA extraction, amplification, and sequencing

One to three legs of each specimen were typically used for total genomic DNA extraction, voucher specimens are deposited in the University of Hawaii Insect Museum (UHIM). Some specimens were additionally used for genomic studies (e.g. (Dupuis et al. 2018a), such flies were fully ground up to obtain higher concentration DNA extracts. DNA was extracted using the Qiagen (Valencia, CA) DNeasy animal blood and tissue kit, following manufacturer’s protocols. A 780 base-pair (bp) section of the Cytochrome C Oxidase I 3’P region (COI-3P) of the mitochondrial DNA was amplified using the forward primer HCO-2198rc (5’-3’ GCT CAA CAA ATC ATA AAG ATA TTG G; (Folmer et al. 1994) and reverse primer Pat-k508 (5’-3’ TCC AAT GCA CTA ATC TGC CAT ATT A; (Simon 1994). The PCR thermal conditions were 2 min. at 94°C, 40 cycles of (94°C for 30 sec., 53°C for 30 sec. and 70°C for 60 sec.) with a final extension at 70°C for 10 min. Bidirectional Sanger sequencing was outsourced to Eurofins (Louisville, Kentucky, USA). We combined chromatograms of the forward and reverse reads in Geneious v.R10, trimmed primer regions, and created alignments for each species. The alignments contained no insertions or deletions, and all sequences were checked for stop-codons, which can indicate the presence of nuclear pseudogenes. The sequences are deposited both in BOLD (DOI:xxx.xxx.xxx) and NCBI Genbank, accession numbers xxxx-xxxx.

### Phylogenetic and population genetic analyses

The final datasets used for analyses included 428 *B. correcta*, 756 bp aligned, 214 *B. zonata*, 752 bp aligned and 372 *Z. cucurbitae*, 764 bp aligned. Some papers have studied the COI diversity of *B. correcta* regionally (Jamnongluk et al. 2003; Kunprom et al. 2015), but because of only partially matching COI target regions (i.e. COI-5P versus COI-3P) we have decided to not include their data in our study. Also, Kunprom et al. (Kunprom et al. 2015) reported a strongly divergent COI lineage within *B. correcta*, matching earlier results based on the COI-3P region (Jamnongluk et al. 2003). They attributed this anomaly to *Wolbachia* bacterial infection, but upon comparison of the published COI-3P sequences with our COI reference dataset (data not shown) we conclude this lineage to match a cluster that includes *B. nigrotibialis, B. nigrifacia* and *B. nigrofemoralis*, which share some of the same hosts as *B. correcta* and can be morphologically similar (Allwood et al. 1999; Drew and Romig 2013). For maximum likelihood tree inference of the relationships of the target species and their closest relatives (based on (San Jose et al. 2018a), we used unique haplotypes only and added *B. nigrotibialis* (Perkins), *Zeugodacus tau* (Walker), *Z. synnephes* (Hendel) and *Z. choristus* (May) to the alignments. Maximum likelihood tree inference was done using RaxML v8 with 50 searches for the best tree and subsequent multiparametric bootstrapping with the extended majority rule parameter to estimate statistical support values. We calculated haplotype networks using the TCS statistical parsimony algorithm (Templeton et al. 1992), employed in the software “TCS” v.1.21 (Clement et al. 2000). The raw output of TCS was visualized in the web implementation of tcsBU (Múrias Dos Santos et al. 2015) and optimized for publication in Adobe Illustrator CC. Haplotype rarefaction curves were estimated in R using the HaploAccum() function from the package SpideR with 1,000 permutations, and we used the chaoHaplo() function from the same package to obtain the non-parametric Chao 1 estimator of total haplotype diversity. The “tools and recipes for evolutionary genetics and genomics” (EggLib; (De Mita and Siol 2012) python library was used to calculate the descriptive statistics for nucleotide diversity π, Tajima’s D, Watterson’s estimator of θ, and Fu’s Fs.

## Results

### COI Phylogeny

We successfully obtained COI sequences of 428 *Bactrocera correcta*, 214 *B. zonata* and 372 *Zeugodacus cucurbitae*. The COI maximum likelihood trees (Fig. 2) based on unique haplotypes, with added outgroup sister taxa, show that each of the study species can reliably be separated from its closest sister species using a monophyly criterion, and that the intraspecific variation does not overlap with the variation between species. All haplotypes are unique to each respective species. The intraspecific variation ranges from 1.32% in *Z. cucurbitae* to 3.06% in *B. zonata* (Table 1), and this variation is distributed across the COI region.

**Table 1.** Species-level population genetic statistics

| | Ntaxa | Nhap | Max.<br>intraspecific<br>P-distance | Variable<br>sites | $\theta_w$ | $\pi$ | Fu’s Fs | Tajima’s<br>D |
| --- | --- | --- | --- | --- | --- | --- | --- | --- |
| *B. zonata* | 210 | 184 | 3.06% | 21.5% | 25.161 | 7.423 | -378.223 | -2.205 |
| *B. correcta* | 428 | 80 | 2.12% | 13.5% | 14.318 | 4.224 | -71.638 | -2.061 |
| *Z. cucurbitae* | 366 | 70 | 1.32% | 8.7% | 11.577 | 0.967 | -142.584 | -2.673 |

**Figure 2:**
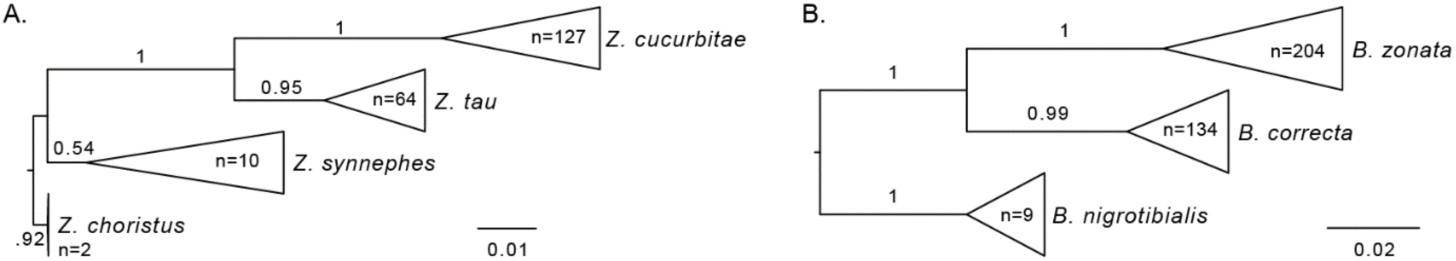
COI maximum likelihood trees based on unique haplotypes showing a monophyletic origin for A. *Zeugodacus cucurbitae* and its closest congeners and B. *Bactrocera zonata* and *B. correcta* and their closest sister species. Scale bar indicates substitutions per site; the taxa are ‘collapsed’ into a triangle for each species, where the horizontal width indicates the maximum intraspecific variation. Bootstrap support values are indicated on the respective branches.

### Haplotype networks

We find contrasting patterns of haplotype diversity between the three species (Fig. 3), where *B. zonata* has the highest diversity and most haplotypes are represented by singletons, *B. correcta* has ∼7 dominantly represented haplotypes and *Z. cucurbitae* has a single dominant haplotype with closely related singeltons in a ‘star-shape’ surrounding it. None of the species displays a biogeographic signal, where certain haplotypes or parts of the haplotype network can be correlated with a biogeographic region. For *B. zonata*, many of the haplotypes are unique to a region, but also unique in the network and further sampling will likely show that these are shared between regions.

**Figure 3.**
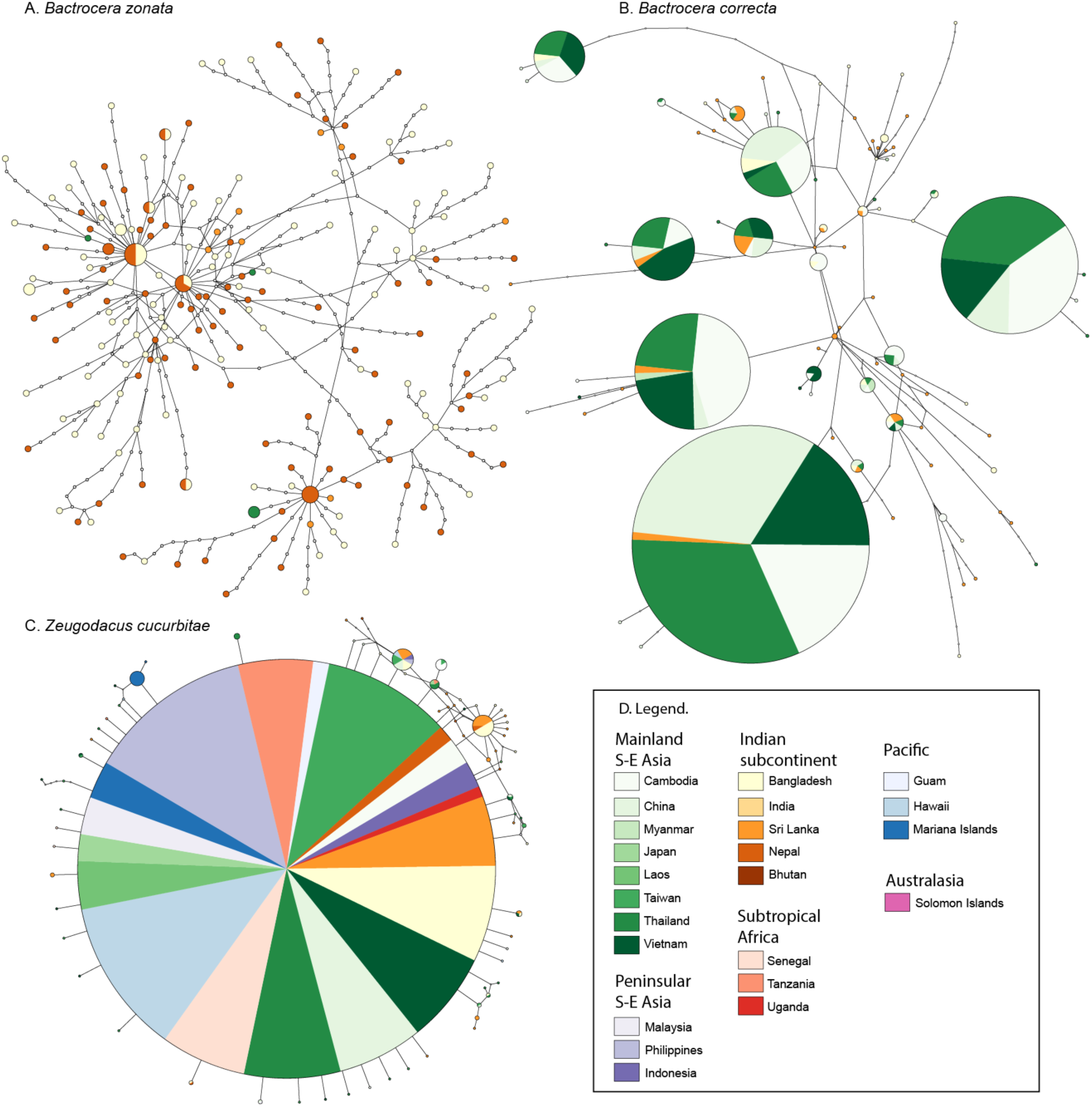
TCS haplotype networks for A. *Bactrocera zonata*, B. *Bactrocera correcta* and C. *Zeugodacus cucurbitae* and the legend for all three in D. Each circle represents a haplotype, circle size indicates the occurrence of each haplotype, but note that the scale differs between A, B and C. The colors refer to the sampling locality, with shades of the same color roughly following biogeographical regions; white circles connecting sampled haplotypes indicate hypothetical (unsampled) haplotypes.

### Haplotype rarefaction

The haplotype rarefaction curves with randomly sampled diversity do not reach an asymptote for any of the species (Fig. 4), indicating that we have only sampled part of the actual diversity. Despite their strongly different haplotype networks (Fig. 3), *Bactrocera correcta* and *Zeugodacus cucurbitae* have largely overlapping haplotype rarefaction curves and confidence intervals, although the Chao 1 estimated total diversity for the former is 1,110 haplotypes and only 270 for the latter. The angle on the curve for *Bactrocera zonata* is close to a 1:1 ratio where every new sample represents a new haplotype, indicating a much higher diversity for this species, which is underlined by a Chao 1 estimated total haplotype diversity of 1,890.

**Figure 4.**
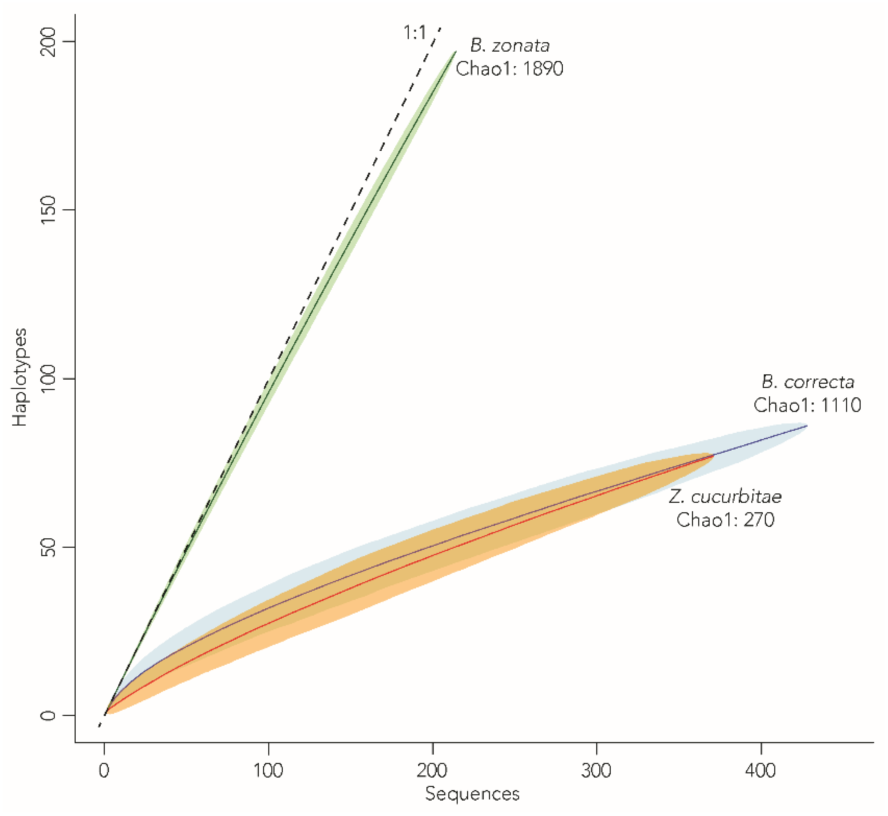
Haplotype rarefaction curves and the Chao 1 estimated total haplotypes for each species, shaded areas indicate 95% confidence interval from 1,000 permutations. The 1:1 line indicates the maximum where each new sample represents a new haplotype.

### Population genetic statistics

From the lack of geographic partitioning in the haplotype networks (Fig. 3), we assumed the different sampling points for each species to be part of a panmictic population and estimated descriptors of their genetics (Table 1). The number of observed haplotypes ranged from 70 to 184. With Chao 1 total estimated haplotypes per species ranging from 270 to 1,890 (Fig. 4.), we sampled 7.2% (in *B. correcta*) – 25.9% (in *Z. cucurbitae*) of the total estimated diversity. Watterson’s estimator of θ and nucleotide diversity π estimators show that *B. zonata* is the most diverse species, followed by *B. correcta*, and *Z. cucurbitae* shows little diversity, matching the results from the haplotype network (Fig. 3). Wattersons’s estimator of θ, calculated through the formula θw = 4N_e_µ can also be used as an estimate of effective population size (N_e_), and shows that *Z. cucurbitae* has the smallest effective population size, followed by *B. correcta* and *B. zonata*. Negative values of Fu’s Fs are evidence for an excess number of alleles relative to the expected. Fu’s Fs is negative for all three species, but the largest negative is in *B. zonata*. An excess of low frequency polymorphisms relative to expectation is expressed by negative values of Tajima’s D, and are commonly interpreted as demographic estimators for population expansion or selection. We found negative values of Tajima’s D for all three species, but most strongly in *Z. cucurbitae*, followed by *B. zonata* and *B. correcta*.

## Discussion

### Distance decay COI diversity

We examined the role of the distance-decay model in the mitochondrial gene COI as a tool to understand the degree of spatially heterogeneity in genetic diversity and whether important pest species have experienced recent range expansions. In general, intraspecific genetic diversity is a result of demographic history, mutation and selection (Bazin et al. 2006). The different genetic patterns observed can reveal much about the forces, such as rapid expansion or significant bottlenecks, directly affecting species of interest. Considering the prevalence of horticulture in human history, it is difficult to assess what should be considered the native range of the three species. The native range can sometimes be assessed a posteriori, because it will commonly have the highest genetic diversity (e.g. Lees et al. 2011), but a priori assumptions rely on historical records of observations and these may be inaccurate due to, e.g., faulty taxonomy (Schutze et al. 2015). For our study, we therefore attempted to include as many samples from throughout the current ranges of the species as possible, and we did not detect variation in haplotype diversity between regions that would enable the distinction between native and non-native ranges. Despite the fact that we did not detect regional variation, all three species in our study show markedly different COI diversity patterns. This might be expected to influence their invasion success if genetic diversity is a boon for invasive species. The differences may be a result of the differing topology of the landscape of their range, ecological interactions and/or species-specific variation in the mutation rate (µ) of COI. *Zeugodacus cucurbitae* has the widest distribution in our study, yet, conversely, it shows a lack of haplotype diversity. This could be due to very recent expansion in *Z. cucurbitae* from a very genetically limited source population that has adapted to the agricultural environments and since spread across the globe. The pattern of a single adapted population spreading rapidly due to humans has also been observed in other organisms (e.g. (Branch and Nina Steffani 2004; Rubinoff et al. 2010; Reil et al. 2018). Regardless of the underlying reasons for the contrasting patterns found in this study, the diverging haplotype patterns suggest that distance decay based principles likely do not apply to these pest species, at least not anymore. As a consequence, the management strategies undertaken for each pest should be based on the absence or presence of intraspecific diversity in regional populations observed. These results also pertain to species of conservation concern, since understanding current and past population retractions as measured by genetic diversity across their range has important implications for the evaluation and identification of declining taxa.

### Invasion history and potential

Inherent to the host preferences of these insects, there is a fairly detailed record of their range expansion (Meyer et al. 2007, 2013; Vargas et al. 2015). The intraspecific mtDNA diversity reveals additional aspects of the invasive history and, possibly potential by assessing the demographic history and biogeographic distribution of diversity. The low levels of variation and star-like shape of the mtDNA genealogy of *Z. cucurbitae* are consistent with a sudden population expansion. Large negative values of Fu’s *F*s and Tajima’s *D*-tests further indicate a population demographic expansion (Schmidt and Pool 2002; Pramual et al. 2011). In stark contrast with these results, we found a new haplotype for almost every additional sample of *B. zonata* that we included. The demographic estimators for the latter do not suggest that there has been rapid population expansion, at least not from a small source population, and in the regions covered by our sampling. A study of the COI variation of 1,600 specimens of two other prominent pests of the tribe, *B. dorsalis* and *B. carambolae,* also showed high levels of haplotype variation and a large global distribution with multiple recently invaded continents (San Jose et al. 2018b). Overall, the results of Tajima’s D estimator seem congruent with the historical record of invasions. *Zeugodacus cucurbitae* has the largest negative value for Tajima’s D and is the most invasive and widespread of the three pests, *B. zonata* is the second and *B. correcta* third. However, it is worth noting that the melon, gourd and squash hosts of *Z. cucurbitae* are also the most widely grown and transported. The demographic estimators can reveal if there have been recent changes in population sizes, but the values depend on the original population structure in the ‘native’ range and only partly reflect the species’ pest ‘potential’. The pest potential, which can be regarded as the combined factors of the likelihood of reaching new areas and the possibility of becoming established (Vargas et al. 2015), is likely mostly determined extrinsically, by human actions – such as the growing of host plants and transporting of fruit. It appears that depending on these circumstances, any of the species in the tribe can develop into pests, with serious implications for quarantine which currently focuses on the handful of already known pest species.

### Undetected(?) biogeographic patterns

Although COI has been shown to reveal geographic patterning in some species (e.g. (O’Loughlin et al. 2008; Valade et al. 2009; Morgan et al. 2010), we did not detect any biogeographic structure in our three pest species. The effects of historical climatic change were not as severe in Southeast Asia as in temperate regions, with more continuous possibilities for gene-flow throughout the region which may have resulted in less strongly pronounced biogeographic patterns (Penny 2001; Cannon and Manos 2003). However, a high resolution (1,097 SNPs) study of the melon fly *Z. cucurbitae* revealed geographic structuring with six to ten clusters that were only partly detected using microsatellite data (Dupuis et al. 2018a), and are not detected in our current study using COI. Despite being a widespread species occurring from Africa across Asia into the Pacific, we only found a single dominant haplotype. This supports the scenario from the demographic estimators and the star-shape of the haplotype network, all suggesting that *Z. cucurbitae* was restricted to a small geographic area and began to expand rapidly relatively recently– an expansion that is still ongoing as more of Africa is invaded (De Meyer et al. 2015). There are, however, some alternative explanations that we cannot exclude based on our present data, such as the presence of *Wolbachia* bacteria that could have influenced the mitochondrial diversity (Smith et al. 2012). Nonetheless, the SNP data supports biogeographic patterning. The differences between mtDNA versus nuclear DNA are partly due to the differences in mutation rates and inheritance, but also to the fact that the method to derive nuclear DNA SNP’s results in a much larger data set of independent sites with a greater ability to resolve small differences between populations. As more SNP data becomes available for more species, it will be interesting to test at which time-scales, or under which evolutionary circumstances, the resolution between mtDNA and SNP data are similar.

In contrast to the data available for *Z. cucurbitae*, the biogeographic structures of *B. zonata* and *B. correcta* have not been studied with high-resolution (i.e. genomic) methods, which may similarly reveal patterning that we did not detect using COI. The peach fruit fly *B. zonata* is mostly found on the Indian subcontinent and is expanding westward into Africa. Despite the peach reference in its common name, it was likely first introduced into Africa with guava in 1993 (Meyer et al. 2007). The hosts of *B. zonata* and *B. correcta* largely overlap and similar biogeographical and population patterns might have been expected. Based on their current distribution, overlapping host range, phylogenetic relation and similar morphology, the most parsimonious conclusion is that *B. correcta* and *B. zonata* speciated through vicariance. *Bactrocera correcta* is now found mostly in eastern Asia, and *B. zonata* is mostly found in the Indian subcontinent with recent westward expansion (De Meyer et al. 2015). The two species only overlap in Sri Lanka, India and Thailand. These vicariant distributions, despite the likely possibilities for human-mediated dispersal (fruit trade) into either range, may indicate strong competition or selection pressures that limit their avenues for invading regions already occupied by the other species. Fragments of the COI barcode region (COI-5P) of *B. zonata* have been studied in India (Choudhary et al. 2017), where they also concluded that *B. zonata* has high intraspecific variability, but showed no biogeographic patterning. The third species in our study, the guava fruit fly *Bactrocera correcta* is mostly found in mainland Southeast Asia. A regional study of the COI-5P diversity of *B. correcta* in Thailand concluded that there was much genetic homogeneity, likely because host plants are spatially and temporally continuously available (Kunprom et al. 2015). However, we found samples from Thailand to belong to at least nine different haplotypes, mostly those dominantly represented in our dataset. All three species pose large invasion risks and biogeographical analysis of intercepted specimens can help to diagnose pathways and block them (e.g. (Shi et al. 2012; San Jose et al. 2018b). However, from the lack of biogeographic patterning we conclude that COI data cannot be used to trace the origins of introductions or intercepted flies of *Z. cucurbitae, B. correcta* and *B. zonata*.

### COI-based identification

Percentage-wise, the ranges for intra- and interspecific COI variation we find in our study species are similar to those found in other groups of life, such as moths, beetles, or birds, which commonly show <3% intraspecific and >2% interspecific variation (Kerr et al. 2007; Rivera and Currie 2009; Pentinsaari et al. 2014; Schmidt et al. 2015; Huemer et al. 2018). Even though the estimated total number of haplotypes between the three species varies seven-fold, the maximum intraspecific diversity is at most ∼3% and all species can be identified reliably using COI. The high mutation rates in COI, although predominantly in the third-codon position and synonyms are generally known across all animal taxa (Pentinsaari et al. 2016), and have made the marker a useful part of a broader dataset for species delimitation and recognition, and to some extent population-level studies. Contrasting patterns of COI evolution between higher groups has been observed before (Meier et al. 2006; Pentinsaari et al. 2016), which has been linked to the functional properties of the gene, but here we show that large differences can exist between closely related (sister) species as well, or species within the same tribe. Although we only studied three species, the patterns of haplotype diversity are starkly contrasting with very low haplotype diversity in *Z. cucurbitae* to very high in *B. zonata*. The estimates of the proportion of the diversity that we sampled range from 7.2% - 25.9%. All three of these species are pests associated with crops and would be expected to have high effective population sizes (N_e_), given the potentially high population density in agricultural systems. The statistic Watterson’s θ is dependent on N_e_ and genetic diversity µ, and it is possible that only µ varies significantly between these three pests. Mitochondrial genetic diversity, in contrast to nuclear genetic diversity, may not be linked to N_e_ but only to µ and selection (Wang 2005; Bazin et al. 2006). In *Bactrocera dorsalis* and *B. carambolae,* high mutation rates were postulated to cause hyper-diversity in their mitochondrial genomes (San Jose et al. 2018b). However, another explanation for this pattern may have to do with the demographic and invasion history of each species. Despite the common application of COI sequence data for large groups of life across the planet (Kerr et al. 2007; Steinke et al. 2009; Huemer et al. 2014; Mutanen et al. 2016), there are many open-ended questions on how this molecule functions, and to what extent there is active selection (Rubinoff et al. 2006; Pentinsaari et al. 2016). There have been few studies thus far on the intraspecific variation of COI using large sample sizes, but as DNA sequencing continues to become cheaper (e.g. (Hebert et al. 2018) there will hopefully be larger sample sizes for more groups and we can evaluate to what extent general rules apply and how many samples are required to cover the haplotype diversity, to ensure reliable species identification.

## Conclusion

Our results indicate that COI was effective for species level identification across the ranges of these three invasive pest species, even in the presence of considerable intraspecific heterogeneity among species. Generalized distance decay predictions may not be applicable for pest species, because recently expanded distributions due to globalization may obscure historic biogeographic patterns. As a result, we could not use COI intraspecific variation to identify source populations or invasion pathways to augment pest management, and more high-resolution genomic methods may be required. We found that simple demographic estimators based on COI haplotype diversity can be used to infer the historical expansion speed and complement otherwise documented distribution data, but that it cannot predict the virulence of a pest because its spread is more likely dependent on extrinsic factors, such as globalization.

## Acknowledgements

We greatly appreciate help from Dan Nitta, Julian Dupuis and Kimberly Morris with the molecular work for efficiently processing large numbers of samples. We would also like to thank Scott Geib for allowing us kind use of his laboratory resources. We further thank Ken Bloem for providing comments that improved the manuscript. Funding for this project was provided by the United States Department of Agriculture (USDA) Farm Bill Section 10007 Plant Pest and Disease Management and Disaster Prevention Program in support of suggestion “Genomic approaches to fruit fly exclusion and pathway analysis”: 3.0497-FY17. These funds were managed as cooperative agreements between USDA Animal and Plant Health Inspection Service and the University of Hawaii’s College of Tropical Agriculture and Human Resources (8130-0565-CA) and the University of Idaho’s College of Agriculture and Life Sciences (8130-0665-CA). This material was made possible, in part, by a Cooperative Agreement from the United States Department of Agriculture’s Animal and Plant Health Inspection Service (APHIS). It may not necessarily express APHIS’ views. Additional funding was provided by the USDA Cooperative State Research, Education and Extension (CSREES) project HAW00942-H administered by the College of Tropical Agriculture and Human Resources, University of Hawaii.

